# Genomic databanks and targeted assays help characterise domestic mosquito incursions

**DOI:** 10.1101/2022.01.19.477022

**Authors:** Thomas L. Schmidt, Nancy Endersby-Harshman, Nina Kurucz, William Pettit, Vicki L Krause, Gerhard Ehlers, Mutizwa Odwell Muzari, Bart J Currie, Ary A Hoffmann

## Abstract

Biosecurity strategies that aim to restrict the spread of invasive pests can benefit from knowing where new incursions have come from. This knowledge can be acquired using genomic databanks, by comparing genetic variation in incursion samples against reference samples. Here we use genomic databanks to investigate domestic incursions of two mosquito species in Australia, and assess the performance of genomic tracing methods when databank samples were collected some time ago or are genetically similar. We used a deep learning method to trace a 2021 invasion of *Aedes aegypti* in Tennant Creek, Northern Territory, to Townsville, Queensland, and to trace two years of *Ae. albopictus* incursions to two specific islands in the Torres Strait. We observed high precision of tracing despite 30-70 generations separating incursion and reference samples. Targeted assays also provided additional information on the origin of the Tennant Creek *Ae. aegypti*, in this case by comparing *Wolbachia* infection data and mitochondrial DNA variation. Patterns of relatedness and inbreeding indicated that Tennant Creek was likely invaded by one family of *Ae. aegypti*, whereas Torres Strait incursions involved distinct kinship groups. Our results highlight the value of genomic databanks that remain informative over years and for a range of biological conditions, and demonstrate how additional targeted assays (e.g. *Wolbachia*) can improve inferences.

**Key Message:**

- Genomic tracing can provide valuable information on pest incursions and new invasions.
- Evolution will lead to increasing differences between databanks and extant populations.
- We tested how well genomic databanks could trace incursions sampled 30-70 generations later and where genetic differentiation was low.
- We show that tracing methods are robust for a wide range of conditions, and report specific incursion origins for two *Aedes* species.
- Our results suggest that genomic databanks will remain informative over years and for a range of invasive systems.

## Introduction

Invasions of exotic pests confer a global economic burden of over $200B annually (Diagne et al. 2021). These costs have risen in recent decades and are expected to rise further as invasions continue to increase in frequency worldwide (Sardain et al. 2019). For invertebrates, most invasions establish via accidental transportation on ships, planes, or land vehicles, and are typically hard to eradicate with insecticides due to resistance that is either present on arrival or evolves shortly afterward (Gao et al. 2021). Border biosecurity operations have demonstrated success at restricting the spread of pests (Caley et al. 2015; Muzari et al. 2017), however, resource limitations constrain the number of inspections on arriving vessels and the number of border sections that can be effectively monitored.

Biosecurity operations can be greatly assisted by knowledge of incursion pathways, as this allows for more frequent inspection of goods and conveyances from likely source locations (Robinson et al. 2011) and regular inspection of known entry points (Myers et al. 2000; Mehta et al. 2007). While pathways may be easily inferred if incursions are detected on their vessel of entry, incursions that are detected later pose a greater challenge. Recent work using genomic databanks to trace incursions or recently established invasions indicates that these methods can reliably infer the most likely origin out of a set of reference populations (Gloria-Soria et al. 2018; Schmidt et al. 2019; Sherpa et al. 2019; Schmidt et al. 2020b; Chen et al. 2021; Kelly et al. 2021; Popa-Báez et al. 2021; Yan et al. 2021; Schmidt et al. 2021c), and are particularly useful for tracing samples collected beyond border biosecurity. These methods compare genome-wide genetic variation in the incursion or invasion of interest against variation across reference samples, allowing the invasion to be linked to a population or a set of related populations of origin depending on the quality of the databank.

This study presents genomic tracing results for two mosquito species, the dengue mosquito *Aedes aegypti* and the Asian tiger mosquito *Ae. albopictus*. Previous work on tracing these mosquitoes in Australia has focused on determining the country of origin of incursion samples collected at airports and seaports (Schmidt et al. 2019, 2020b), and some tracing of *Ae. albopictus* has also been completed within the Torres Strait Islands, Queensland, as part of a study considering population structure across this region (Schmidt et al. 2021c). Here we focus on new in-country detections within Australia away from entry ports where invasive samples of unknown provenance have been detected. For *Ae. aegypti*, we trace an invasion of Tennant Creek, Northern Territory, that was detected in early 2021. For *Ae. albopictus*, we trace a set of new incursions detected in the Torres Strait on islands where this species has been successfully eliminated (Muzari et al. 2017).

While pest management is beginning to employ genomic tracing, studies have yet to investigate the tracing power of genomic databanks or how tracing fares after tens of generations of drift and migration. These questions are vital for the wider uptake of pest genomics as they determine the necessary size of the initial sequencing investment and the length of time a databank will remain informative for tracing. We show here that tracing with the deep learning method Locator (Battey et al. 2020) is remarkably robust, as seen in high levels of unambiguous assignment during cross-validation of the reference databanks, and in the precise tracing of new incursions despite the age of the *Ae. aegypti* databank samples and high genetic similarity in the *Ae. albopictus* samples (Maynard et al. 2017; Schmidt et al. 2021c). We also show how tracing precision was improved by targeted assays for insecticide resistance and a *Wolbachia* infection, which follow releases to establish *Wolbachia* in north-eastern Australia over the last decade (Schmidt et al. 2017; O’Neill et al. 2018; Ryan et al. 2020). Finally, we show how genetic relatedness can be used to infer other demographic parameters, in this case that the *Ae. aegypti* invasion was likely sourced from a single geographical location and potentially from a single family while the *Ae. albopictus* incursions were from different family groups.

## Materials and Methods

### Mosquito sample collection

#### *Aedes aegypti* invasion of Tennant Creek, NT, Australia

*Aedes aegypti* was detected in Tennant Creek during routine surveillance between 22^nd^ – 26^th^ February 2021. Four adult mosquitoes from this new invasion (three females, one male) were collected from a single property at 26^th^ February. Follow-up sampling between 9^th^ - 11^th^ March located an additional five adults (three females, two males) and nine immatures at four properties within ~80 m of the original detection.

To trace the source of this invasion, we built a genomic databank from 80 *Ae. aegypti* sampled from 11 locations across its Australian distribution in Queensland, from the northern islands of the Torres Strait (9.3° S) to Goomeri in southeast Queensland (26.2° S) (Figure 1a). These were sampled between 2009 and 2017. We also included previously-generated sequence data from Cairns (Schmidt et al. 2018) and Townsville (Rašić et al. 2016), sampled in 2015 and 2014 respectively. The Cairns sample included some individuals infected with *Wolbachia* from earlier releases (Schmidt et al. 2017) while the Townsville sample was taken before *Wolbachia* releases in this area (O’Neill et al. 2018).

**Fig. 1.**
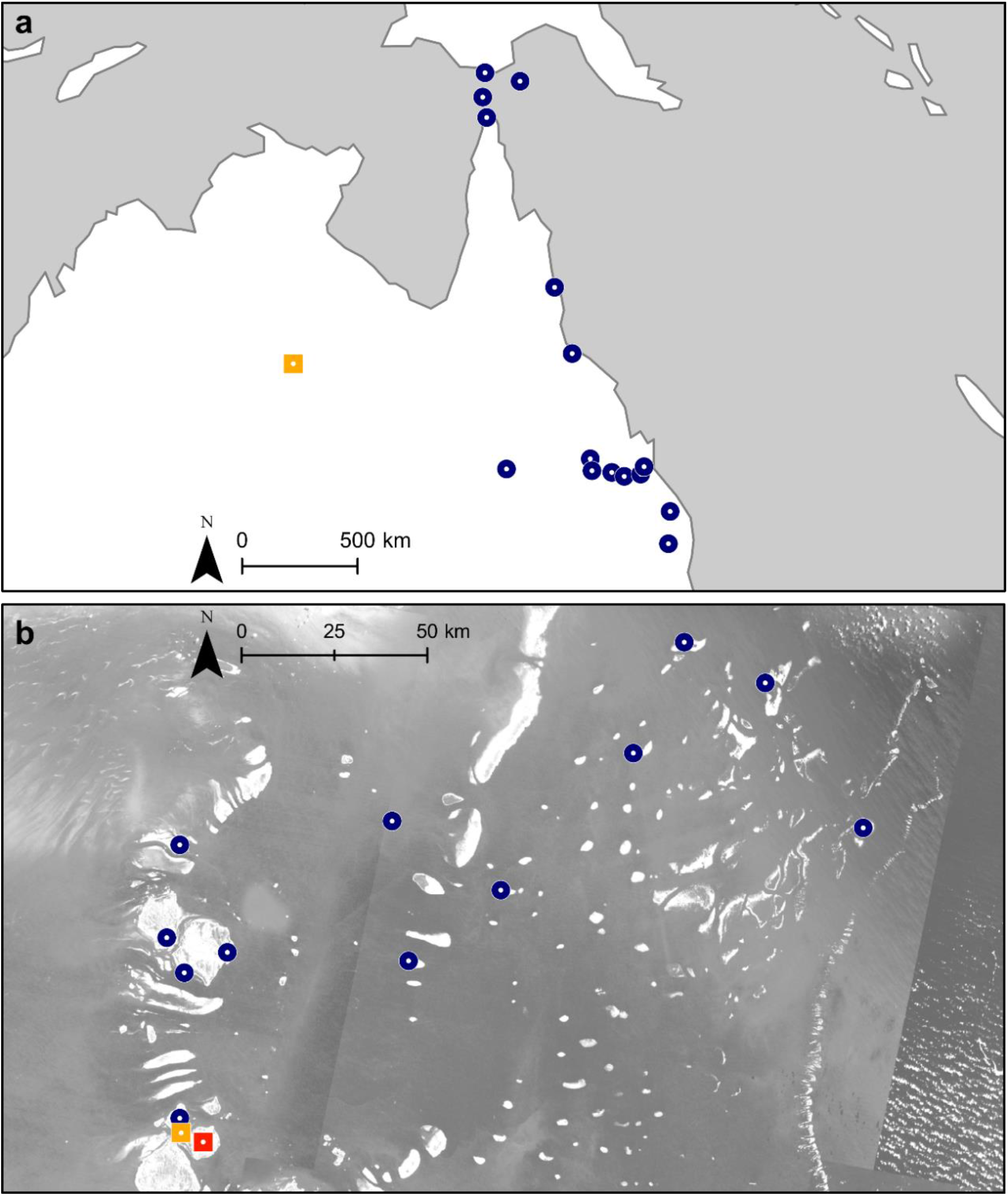
Locations of *Ae. aegypti* (a) and *Ae. albopictus* (b) samples. Blue circles show locations of reference samples. In (a), the orange square indicates Tennant Creek. In (b), the orange square indicates Thursday Island and the red square indicates Horn Island.

#### *Aedes albopictus* incursions in the Torres Strait Islands, QLD, Australia

*Aedes albopictus* incursions were detected during routine surveillance on two islands in the Torres Strait where the species has been successfully eliminated (Muzari et al. 2017). Surveillance conducted in January and February 2021 detected four adult individuals from Thursday Island and three from Horn Island (Figure 1b). Surveillance in January 2022 detected a further four adults on Thursday Island and five on Horn Island. In both years and on both islands, incursions were detected across a wide geographical range (Figures S1-4). There were no further detections of *Ae. albopictus* on either island over the periods March-December 2021 and February-June 2022, suggesting that *Ae. albopictus* had not become established on these islands, and that each detection most probably represented a relatively recent incursion.

To trace the source of this invasion, we used a genomic databank containing previously generated sequence data of 240 *Ae. albopictus* from 12 Torres Strait Island communities across 11 islands, collected in 2017 (Schmidt et al. 2021c) (Figure 1b). Before tracing among Torres Strait Island locations, we checked to ensure that the sample was not from either Papua New Guinea or Indonesia using the same methods as for other tracing and found that they all had a Torres Strait Island origin.

### DNA sequencing and bioinformatic processing

Genomic DNA was extracted from the 98 *Ae. aegypti* and 7 *Ae. albopictus* using Qiagen DNeasy^®^ Blood & Tissue Kits (Qiagen, Hilden, Germany). Extracted DNA was used to build double digest restriction-site associated DNA (ddRAD; Peterson et al. (2012)) sequencing libraries following the protocol of Rašić, Filipović, Weeks, & Hoffmann (2014). A technical replicate was produced of one of the Torres Strait Island incursion samples from Thursday Island, TI-1, which was replicated through the entire protocol after DNA extraction. Libraries were individually barcoded, then sequenced on a HiSeq 4000 using 150 bp chemistry.

New and old sequence data were combined for each species, and run through the same bioinformatic pipeline. We used the Stacks v2.54 (Rochette et al. 2019) program process_radtags to demultiplex sequences, trim them to 140 bp and remove sequences with Phred scores below 20. As the Cairns and Townsville libraries were produced using 100 bp chemistry, we trimmed these to 80 bp. Sequences were aligned to the nuclear genome assembly for *Ae. aegypti*, AaegL5 (Matthews et al. 2018), and the linkage-based assembly for *Ae. albopictus*, AalbF3 (Boyle et al. 2021). Alignment was in Bowtie2 v2.3.4.3 (Langmead and Salzberg 2012) using “–very-sensitive” settings. Sequences were built into catalogs for each species using the Stacks program ref_map. The Stacks program “populations” was used to export VCF files containing SNP genotypes for all individuals in each catalog, filtering to retain SNPs called in at least 50% of individuals from each population and 90% of individuals total, and with a minor allele count ≥ 3 (Linck and Battey 2019). Two of the 18 Tennant Creek *Ae. aegypti* and 1 incursive *Ae. albopictus* from Thursday Island in 2022 had too much missing data for genomic analysis and were omitted, along with 10 *Ae. aegypti* and 3 *Ae. albopictus* reference samples.

### Cross-validation and filtering of reference datasets

To assess the robustness of the reference datasets we performed cross-validation. Each individual in turn was omitted from the training set and treated as an individual of unknown location, and then Locator (Battey et al. 2020; see below) was used to infer the geographical origin of the individual. Assignments were assessed qualitatively for accuracy (i.e. is its origin unambiguous) as well as quantitatively for precision (i.e. how far from its true origin). For the *Ae. aegypti* dataset, we treated nearby samples from Bluff, Capella, and Emerald (QLD) as one location, as well as those from Duaringa, Mt Morgan and Rockhampton (QLD) and those from Gin Gin and Goomeri. This was because sample sizes were low (5) from each specific location. Cross-validation was also used to test two site filtering protocols: where SNPs were not filtered for linkage; and where SNPs within 25 kbp of other SNPs were filtered.

### Incursion tracing

#### i) Genome-wide genetic variation

We used Locator (Battey et al. 2020) to infer the origin of the 16 invasive *Ae. aegypti* and the 16 incursive *Ae. albopictus* (including the technical replicate). Locator is a deep learning method that uses individual genotypes of known geographical location (i.e. the reference individuals) to predict the locations of a set of individuals of unknown location (i.e. the incursive mosquitoes). For each incursive mosquito, we set Locator to provide a point estimate of its origin and 1,000 bootstrap subsamples to indicate confidence around these estimates.

#### ii) *Wolbachia* infection

The Tennant Creek *Ae. aegypti* were assessed for infection of *w*Mel *Wolbachia*, which are present at high frequency in *Ae. aegypti* populations in Cairns, Townsville, and several other locations in northern Queensland, but not in central Queensland populations. *Wolbachia* infection was assessed using a high-throughput PCR assay (Lee et al. 2012) which was run twice on all 18 samples.

In the case of uninfected samples, we also considered the possibility that the *w*Mel infection was initially present but then lost from the invasive population due to the vulnerability of this *Wolbachia* strain to high temperatures (Ross et al. 2017) that are common in Tennant Creek. As *Wolbachia* is coinherited with mtDNA through the maternal line (Hoffmann and Turelli 1997), *w*Mel-infected individuals will all have inherited the same mtDNA haplotype from the same common ancestor within the past ~10 years. This mtDNA haplotype will be retained even if *Wolbachia* has been lost (Yeap et al. 2016), and its presence would be revealed by high genetic similarity in mtDNA between the Tennant Creek *Ae. aegypti* and samples infected with *w*Mel.

To test this hypothesis, we looked at mtDNA variation among our *Ae. aegypti* samples alongside variation from 38 *w*Mel-infected (Wol+) and 87 uninfected wildtype (Wol-) individuals from Cairns, sampled in 2015 (Schmidt et al. 2018). We aligned demultiplexed sequences to the *Ae. aegypti* mtDNA genome assembly (Behura et al. 2011) and called mtDNA SNPs using Stacks. We used VCFtools to output mtDNA genotypes for all individuals with minimum depth of 5, omitting SNPs with any heterozygous genotypes which could represent NUMTs (Hlaing et al. 2009).

#### iii) Pyrethroid resistance mutations

The Tennant Creek *Ae. aegypti* were assessed for three pyrethroid resistance mutations common to *Ae. aegypti* across the Indo-Pacific (Endersby-Harshman et al. 2020), but not within Australia (Endersby-Harshman et al. 2017). For each individual, we ran three replicates of the Custom TaqMan^®^ SNP Genotyping Assays (Life Technologies Corporation, Carlsbad, CA, USA) described in Endersby-Harshman et al. (2020), using on LightCycler II 480 (Roche, Basel, Switzerland) real time PCR machine in a 384-well format.

### Genetic relatedness and inbreeding in the incursions

We investigated patterns of genetic relatedness (*k*) among pairs of individuals (dyads) in the Tennant Creek and Torres Strait incursions, and calculated Wright’s inbreeding coefficient (*F*) for every individual. The patterns can help indicate whether the samples at either location had likely arrived as part of a single incursion or several distinct incursions. For instance, if several incursive individuals had arrived via the same incursion on the same vessel, dyads of these individuals should have higher *k* scores than other dyads in the sample, as they will either be from the same family or have high genetic similarity due to their shared geographical origin. Alternatively, if the invasion was sourced from a single family and related individuals have subsequently mated together, we would expect to see higher *F* scores than in other populations. If neither of these patterns are present, it would suggest that invasive samples represent different incursions.

We used Loiselle’s *k* (Loiselle et al. 1995) to estimate genetic relatedness, calculated in SPAGeDi (Hardy and Vekemans 2002) using the linkage-thinned datasets for each species. Loiselle’s *k* describes correlations in the frequencies of homologous alleles, summarised across the genome (Loiselle et al. 1995). We used this estimator as it makes no assumptions regarding *F* and is suitable for markers with low polymorphism (Vekemans and Hardy 2004). We calculated *F* in VCFtools (Danecek et al. 2011). As estimators of individual-level variation can be biased by uneven sample size and missing data (Schmidt et al. 2021b), we subsampled and refiltered each databank before calculating *F*. For *Ae. aegypti*, we used the Tennant Creek (n=16), Townsville (n=16) and Cairns (n=15) samples. For *Ae. albopictus*, we used n=15 samples from each reference population and the n=15 incursion samples. We filtered to omit all sites with any missing genotypes (--max-missing 1) and to keep only sites with mean depth ≥ 20 (--min-meanDP 20), and did not filter singletons and doubletons.

## Results

### Cross-validation and filtering of reference datasets

Cross-validation indicated that different filtering setting were optimal for each species (Figure 2). In *Ae. aegypti*, 5 reference samples were assigned to the incorrect origin when SNPs were thinned for linkage, and 7 without thinning. Unambiguous assignment of a sample to its true origin occurred in 90 samples (thinned) or 78 samples (no thinning). The average error was 47.1 km (thinned) or 60.7 km (no thinning), out of a maximum distance between samples of 1881.9 km.

**Fig. 2.**
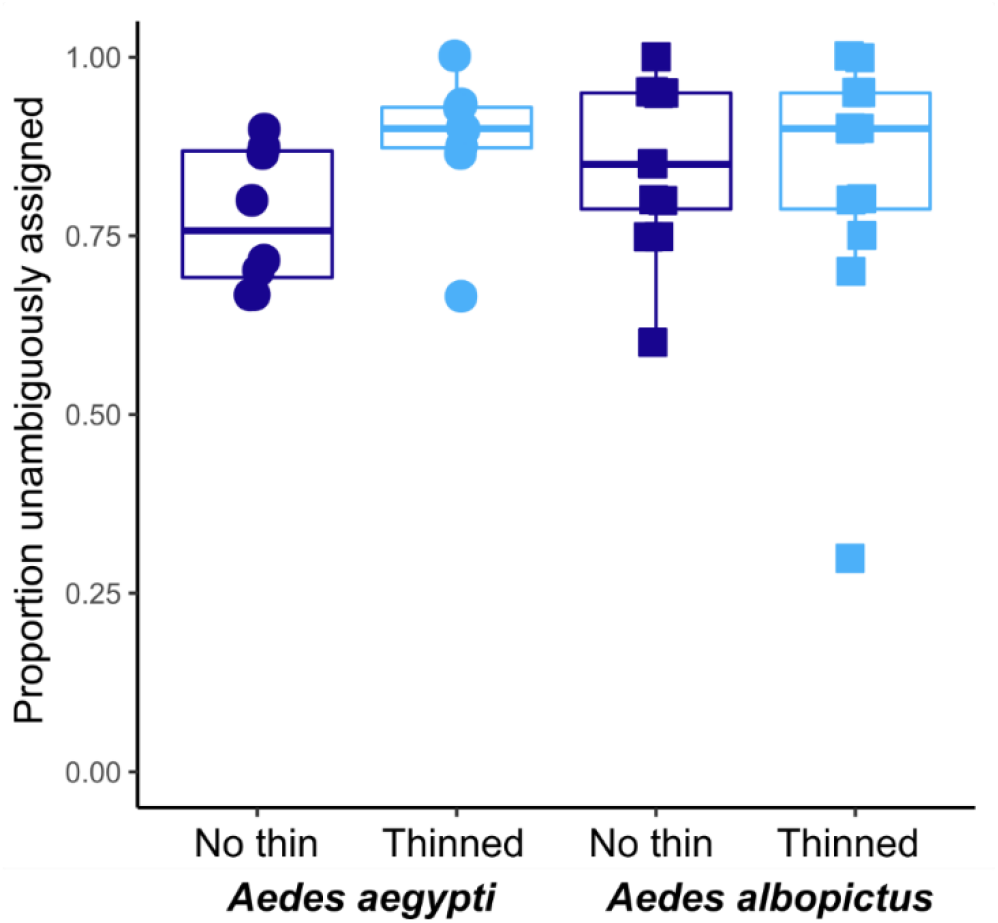
Cross-validation of genomic reference data. SNPs were either not thinned by linkage (dark symbols) or thinned to 25 kbp (light symbols). Points represent the proportion of individuals unambiguously assigned to each reference location.

In *Ae. albopictus*, 10.7% of samples were assigned to the incorrect origin for both thinning treatments. Unambiguous assignment occurred in 199 samples (thinned) or 204 samples (no thinning), and one location, Badu Island, performed particularly poorly in the thinning treatment, with only 30% of individuals unambiguously assigned compared to 60% without thinning. Average error was 11.6 km (thinned) or 11.2 km (no thinning), out of a maximum distance of 213.4 km. There was no evidence in either species that incorrect assignments were biased to specific locations. We used the thinned dataset for *Ae. aegypti* tracing and the unthinned dataset for *Ae. albopictus* tracing.

### Incursion tracing

#### *Aedes aegypti* invasion of Tennant Creek, NT, Australia

Genomic tracing with Locator was run sequentially on different subsets of samples. An initial run with all 102 reference samples and 11866 thinned SNPs indicated all 16 samples had a likely origin between Cairns (16.9° S) and Rockhampton (23.4° S) (Figure S5). Rerunning Locator with reference samples outside this range omitted and 11495 SNPs indicated a likely origin of Townsville (Figure 3).

**Fig. 3.**
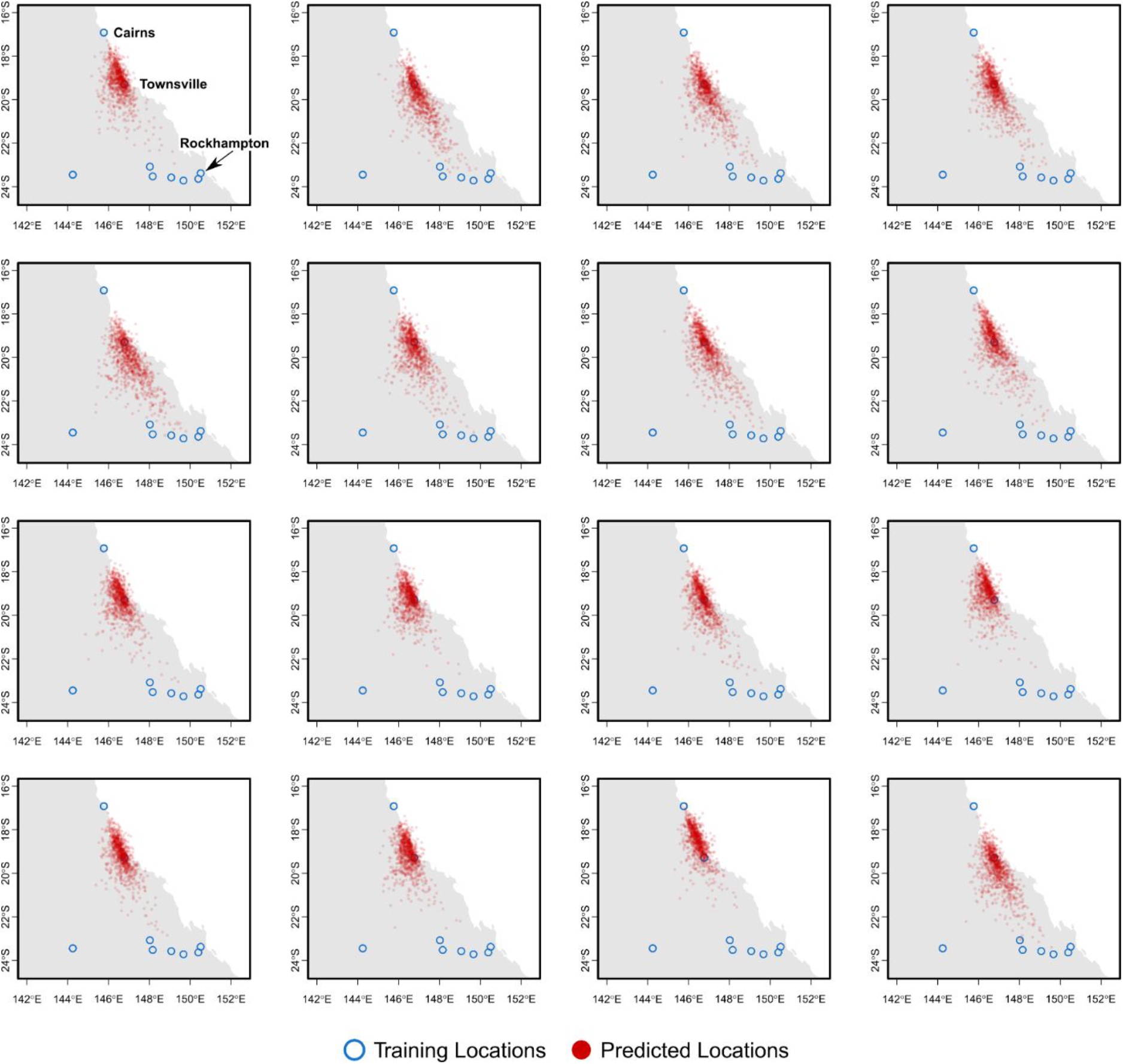
Origins of the 16 Tennant Creek individuals inferred from genome-wide genetic variation. Blue circles indicate reference sample locations, red circles show inferred origins from 1000 bootstrap replicate runs of Locator. Figures S5-7 show results of similar analyses using the whole dataset (S5) or different subsets of the dataset (S6-7)

We checked whether this assignment to origin was biased by Townsville’s geographically central location by rerunning Locator using either Townsville and Cairns only (Figure S6) or Townsville and the southern samples only (Figure S7). In each case, incursives were still placed around Townsville, but Townsville and Cairns were slightly harder to differentiate.

Assays for *w*Mel *Wolbachia* indicated none of the Tennant Creek samples were infected with *w*Mel, in contrast to positive controls from an infected line. Analysis of 116 mitochondrial SNPs showed that pairwise *F*_ST_ between Tennant Creek and Townsville was lower (*F*_ST_ = 0.24) than between Tennant Creek and either the *w*Mel infected or uninfected Cairns samples (both *F*_ST_ = 0.85) (Table 1). This indicates that no *w*Mel infection was ever present in the Tennant Creek samples or their maternal ancestors, otherwise the Tennant Creek sample should be more similar to the Cairns samples given that Cairns provided the genetic background for the original wMel infected strain (Walker et al. 2011). In the pyrethroid resistance assays none of the three resistance alleles was present in the Tennant Creek samples, consistent with the absence of resistance alleles in North Queensland populations (Endersby-Harshman et al. 2017).

**Table 1.**
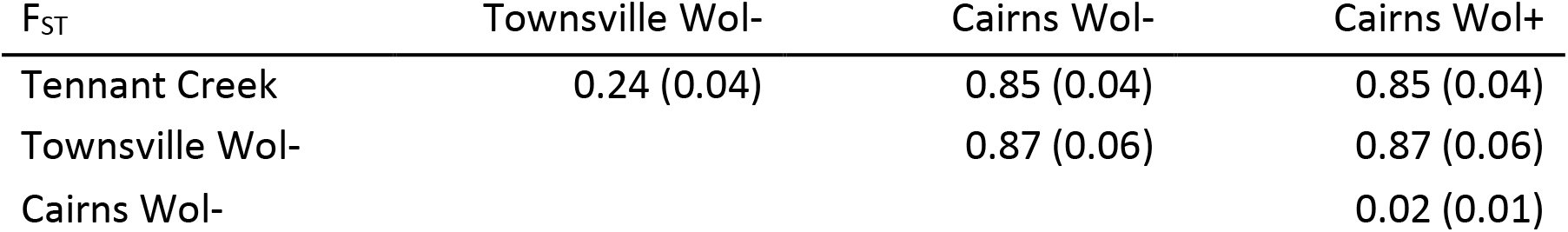
Pairwise *F*_ST_ of mtDNA variation among Tennant Creek, Cairns, and Townsville populations. Wol+ indicates *Wolbachia-infected* samples while Wol-are uninfected. Standard errors are in parentheses.

| $F_{ST}$ | Townsville Wol- | Cairns Wol- | Cairns Wol+ |
| --- | --- | --- | --- |
| Tennant Creek | 0.24 (0.04) | 0.85 (0.04) | 0.85 (0.04) |
| Townsville Wol- |  | 0.87 (0.06) | 0.87 (0.06) |
| Cairns Wol- |  |  | 0.02 (0.01) |

#### *Aedes albopictus* incursions in the Torres Strait Islands, QLD, Australia

Locator was run on *Ae. albopictus* using 73,685 SNPs. Differences in tracing outcomes were evident between the 2021 and 2022 incursions (Figure 4), and these were not due to missing data which was similarly low in both years. For the 2021 samples, the only incursive that could be assigned to a specific location at high confidence was of the Thursday Island sample TI21-1 and its technical replicate, assigned to the St Pauls community on Moa Island. Several of the other 2021 incursives had a likely origin in the Maluilgal (Western) island group, though there was considerable variation among bootstrap replicates. The 2022 samples showed much better assignment, with seven of the nine clearly linked to Keriri Island and six of these showing very low variation among bootstrap replicates. This included one incursive collected next to the airport (HI22-4). Keriri and Moa Island are the two islands to which incursions have previously been linked in this system (Schmidt et al. 2021c).

**Fig. 4.**
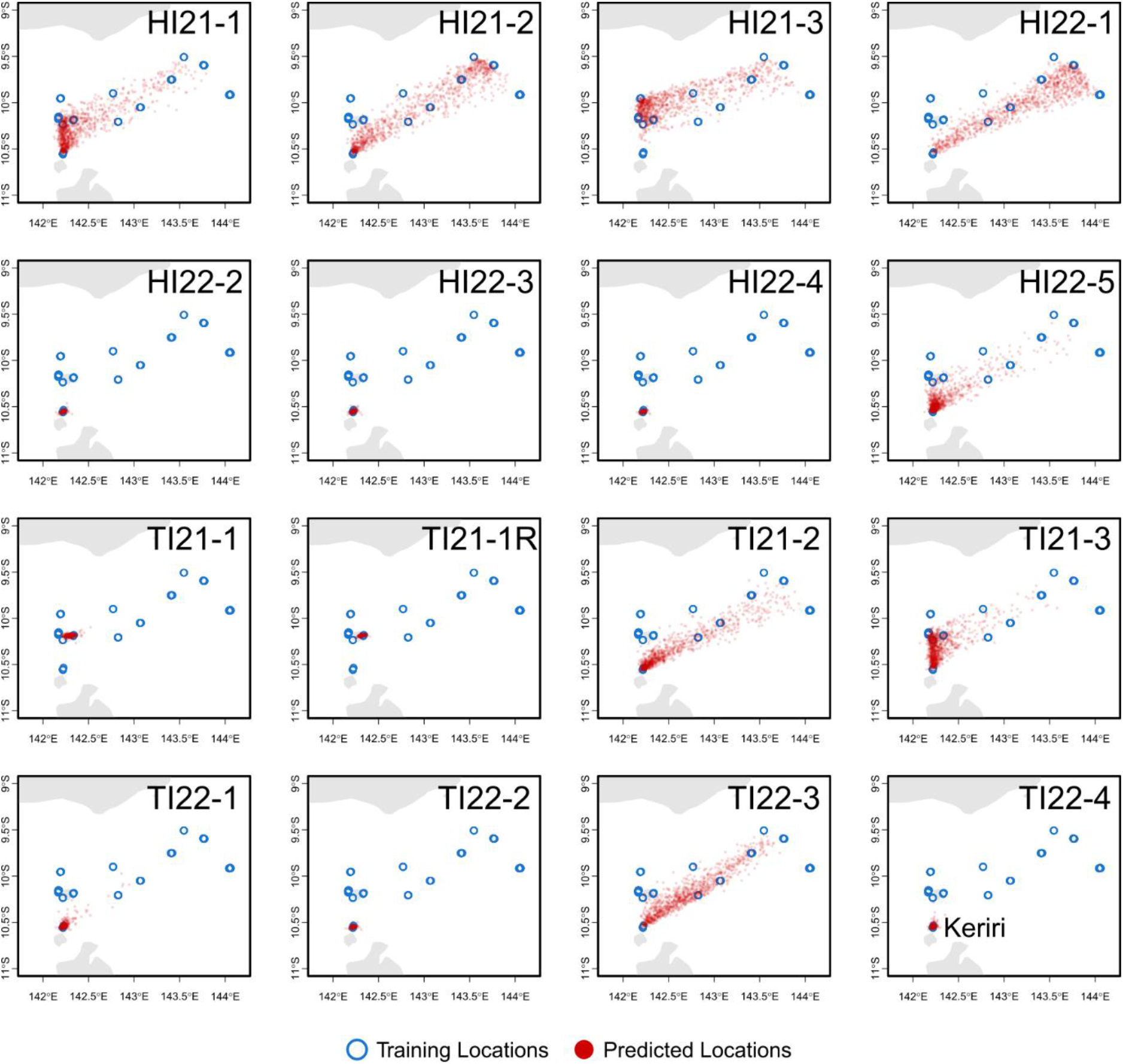
Origins of the 15 Torres Strait Islands individuals inferred from genome-wide genetic variation. Samples were detected on Horn Island (HI) or Thursday Island (TI) in either 2021 or 2022. A technical replicate (R) was made of the TI21-1 sample. Blue circles indicate reference sample locations, red circles show inferred origins from 1000 bootstrap replicate runs of Locator.

### Genetic relatedness and inbreeding in the incursions

Figure 5 displays pairwise Loiselle’s *k* scores for the Tennant Creek and Torres Strait Island dyads in rank order, and Wright’s inbreeding coefficient (*F*) scores in the subsampled and refiltered datasets (5073 SNPs for *Ae. aegypti;* 3223 SNPs for *Ae. albopictus*).

**Fig. 5.**
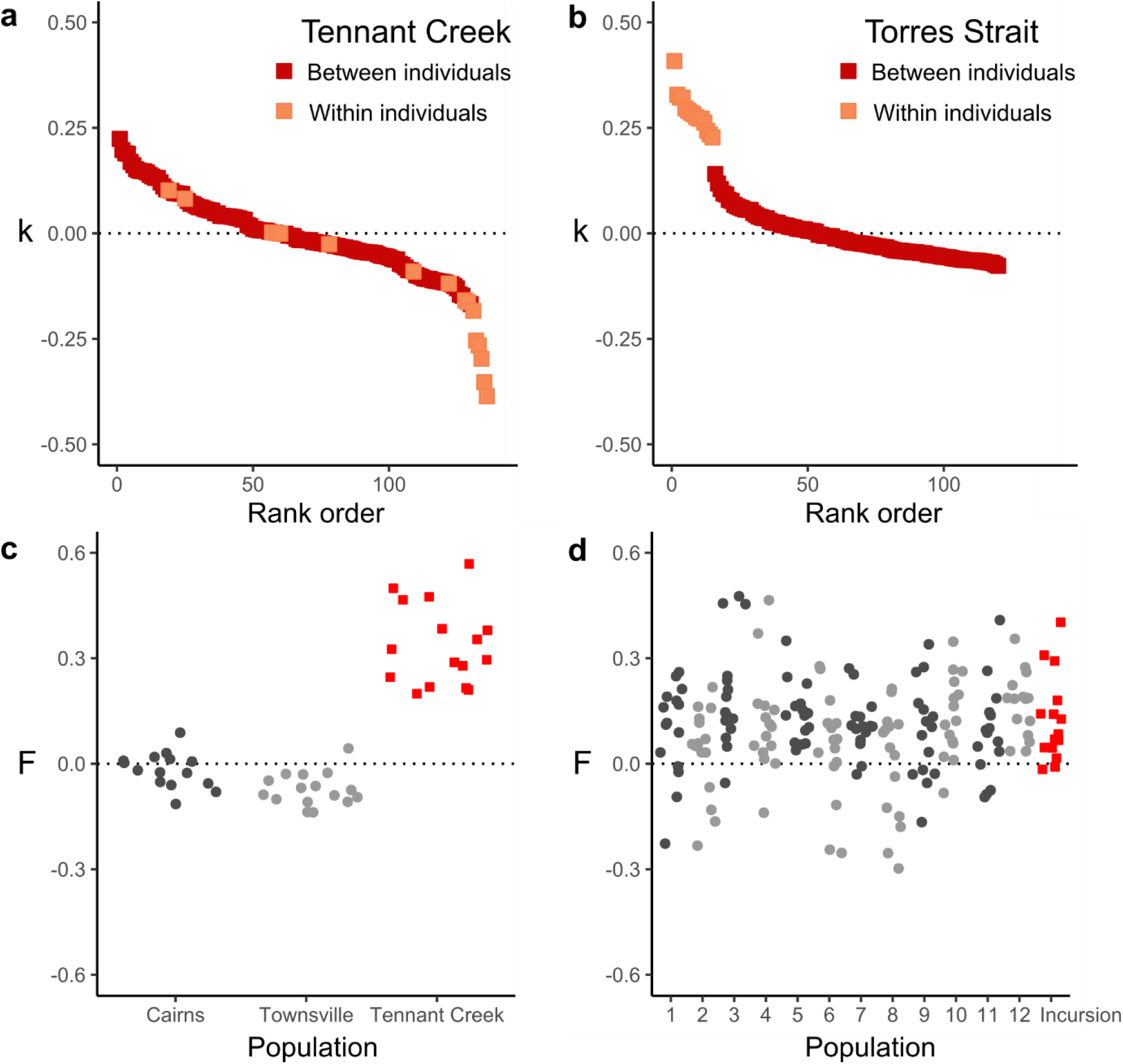
Genetic relatedness (*k*) and Wright’s inbreeding coefficients (*F*) for *Ae. aegypti* (a,c) and *Ae. albopictus* (b,d). Loiselle’s *k* scores are of dyads in Tennant Creek (a) and the Torres Strait Islands (b), where dark squares show correlations between homologous alleles in two different individuals and light squares show correlations within the same individual. *F* scores are of *Ae. aegypti* (c) and *Ae. albopictus* (d), where red squares indicate invasive individuals and grey circles indicate other individuals.

Loiselle’s *k* scores represent either correlations between homologous alleles in two different individuals (dark squares) or the same individual (light squares). The relative magnitudes of *k* scores within and between individuals were different in the two datasets. In the Torres Strait Islands, *k* scores within individuals were all higher than the *k* scores between individuals (Figure 5b). This pattern means that the two chromosomes within each individual are more genetically similar to each other than to chromosomes found in other individuals. The higher *k* within individuals suggests that none of the Torres Strait Island incursives are likely to be from the same family, as dyads from the same family would have chromosomes with high genetic similarity via identity by descent. Instead, these results are evidence that the 15 Torres Strait Island incursives each represent distinct incursions. This pattern was not observed at Tennant Creek, where *k* scores within individuals were mostly undifferentiated from and in many cases lower than *k* scores between individuals (Figure 5a). Chromosomes within individuals are thus no more genetically similar than they are to chromosomes in other individuals. This suggests that the 16 Tennant Creek individuals are all likely to be from the same exact geographical location and are potentially all from the same family.

The above inferences were further supported by variation in *F* scores. In Tennant Creek, *F* scores indicated much higher inbreeding relative to the Cairns and Townsville reference individuals, suggesting that the Tennant Creek invasion was sourced from a single family (Figure 5c, red squares). In the Torres Strait Islands, *F* scores in incursive individuals (Figure 5d) were of similar magnitude to reference individuals, suggesting no inbreeding in incursions.

## Discussion

Pest genomic databanks provide an increasingly popular means of investigating invasions and incursions (Gloria-Soria et al. 2018; Schmidt et al. 2019; Sherpa et al. 2019; Schmidt et al. 2020b; Chen et al. 2021; Kelly et al. 2021; Popa-Báez et al. 2021; Schmidt et al. 2021c). The value provided by genomic databanks should continue to rise as costs of generating genome-scale data from large numbers of pests continue to decrease and as new genomic methods appropriate for pests are developed (North et al. 2021). This study has used genomic databanks to investigate new threats to biosecurity from Australian incursions of the mosquitoes *Ae. aegypti* and *Ae. albopictus*. We were able to infer that the *Ae. aegypti* invasion of Tennant Creek, Northern Territory came from a location near or within Townsville, Queensland that did not have a *Wolbachia* infection at fixation. Eight of the 15 *Ae. albopictus* incursions into the southern Torres Strait Islands were traced to a specific island (Keriri) with very high confidence, with another traced to a second island community (St Pauls on Moa Island). Our estimates of genetic relatedness and inbreeding also showed that the Tennant Creek invasion was likely sourced from one specific location and possibly from a single family, whereas the Torres Strait incursions were all genetically distinct.

As well as answering these specific biosecurity questions, this study helps clarify some broader ideas around the use of genomic databanks in pest management. Most importantly, we have shown via cross-validation that the databanks used in this study are sufficiently powered for precise and unbiased tracing, and our incursion tracing results confirm that this power remains high despite tens of generations of evolution and low geographical genetic structure. If sequences that adequately represent an individual’s true population are not included in the reference dataset, Locator tends to place the individual around the centre of the sampling range (Battey et al. 2020). This was not observed here, as the Tennant Creek invasion was traced to Townsville irrespective of its relative position in the sampling range (Figures S6-7) and the Torres Strait incursives were traced to specific islands in the Western edge of the range (Figure 4).

Cross-validation also indicated different optimal filtering settings for the two databanks. While Locator’s developers have suggested to subsample bootstrap replicates from a set of unlinked SNPs (Battey et al. 2020), we found that thinning led to slightly worse cross-validation in *Ae. albopictus*, though the differences were less pronounced than in *Ae. aegypti* where the unthinned dataset performed much worse (Figure 2). This result may be due to the lower variation in the *Ae. albopictus* databank, which when thinned was reduced to only 4,147 SNPs, as opposed to 14,648 for *Ae. aegypti*. It is currently unclear whether whole genome databanks would provide better results given the larger sequencing effort required for these may have to be offset by smaller sample sizes. Thankfully, whole genome data can be downsampled for compatibility with reduced-representation databanks, which is unfortunately not the case for reduced-representation data produced with different restriction enzymes which currently prohibits analysis of different global datasets (Schmidt et al. 2021a).

In order for databanks to provide value for money, they should be reusable for tracing over years or decades. However, genetic drift and dispersal will ensure that reference samples in the databank are increasingly less representative of extant populations with each passing generation. Our analysis of *Ae. aegypti* showed that the 2021 Tennant Creek invasion samples could be traced to the Townsville genetic background represented by samples collected in January 2014. Assuming 10 generations/year for *Aedes*, the Tennant Creek and Townsville samples should be separated by ~70 generations of evolution, plus several generations of strong genetic drift between invasion and detection. The databank of Torres Strait *Ae. albopictus* showed similar robustness in tracing, wherein a majority of the incursives detected 3-4 years (~30-40 generations) after the reference samples could be traced to two specific islands with very high confidence (Figure 4). This is particularly noteworthy given the much finer spatial scale of the Torres Strait invasion and the low genetic differentiation among islands (F_ST_ ≈ 0.03, Schmidt et al. (2021c)). Differentiation between Cairns and Townsville *Ae. aegypti* is similarly very low (F_ST_’ = 0.018, Schmidt et al. (2020a)) but assignments were unambiguous (Figures 3, S6).

Using targeted assays for *Wolbachia* infection and pyrethroid resistance, combined with mtDNA data from the databank, we were able to increase the confidence of the origin of Tennant Creek *Ae. aegypti* over using nuclear genomic data alone. Importantly, these findings indicate that the invasion may have been sourced from an unsampled location near to Townsville or from a region of outer Townsville where the *w*Mel infection is uncommon, and also that the invasion was unlikely to have come through overseas routes through which resistance alleles are commonly transmitted (Endersby-Harshman et al. 2020). These assays also indicate that the invasion may pose a greater biosecurity threat than otherwise, as without the *w*Mel infection, the Tennant Creek *Ae. aegypti* should be able to transmit dengue without the transmission blocking effects of *w*Mel (Walker et al. 2011).

The different patterns of genetic relatedness observed at Tennant Creek and the Torres Strait provide a useful indication of the genetic characteristics of each invasive system, in that we can infer the incursions into the southern Torres Strait Islands are independent from each other, whereas the invasion of Tennant Creek is from a single source. These genetic inferences align with detection locations, where the Tennant Creek samples were collected within ~80 m of each other while the Horn and Thursday Island incursions were spread across each island’s range. Understanding the independence of incursions can be useful for building predictive models of pest incursion risk as it can indicate propagule pressure (Camac et al. 2021), though these estimates will reflect effective rather than census population sizes of incursions. The findings at Tennant Creek indicate that greater caution will be required for studies seeking to investigate new invasions using close kin dyad methodologies (Jasper et al. 2022; Waples and Feutry 2022). While close kin dyad methods have shown promise for investigating invasions of ~100 generations age (Schmidt et al. 2021c), close kin methods should be applied cautiously to invasions that are very new or that are sourced from a small genetic pool.

## Conclusion

The recent and ongoing drop in sequencing costs has made genomic studies of pests increasingly affordable. A pest genomic databank contains a wealth of information relevant to control, some of which is immediately available (e.g. population structure) and some of which can be revealed over time, such as through tracking the evolution of insecticide resistance or tracing ongoing invasions, incursions, and migration between invaded regions. A central concern of pest genomic tracing is that drift and migration will make databank samples obsolete, requiring frequent resampling and sequencing. This study has investigated these concerns, and found that genomic databanks can remain highly powered for precise tracing for at least 70 generations of evolution. Additionally, cross-validation showed that ~87% of reference samples could be assigned unambiguously, even though both databanks had low genetic differentiation due to proximity or recent invasion histories. These results suggest that pest genomic databanks can provide valuable tracing insights for many years, even at finer geographical scales. Considering our results specifically, in the Torres Strait, the lack of inbreeding among incursives and the predictability of their origins suggest that the current *cordon-sanitaire* biosecurity approach is working effectively, as there is no evidence of cryptic establishment on either island. In Tennant Creek, our results highlight the risks of ongoing reestablishment of *Ae. aegypti* given this invasion was sourced from only a very small number of mosquitoes.

## Author Contributions

All authors conceived and designed research. NK, WP, VLK, GE, OMM and NEH performed sampling. NEH and TLS generated genetic data. TLS analysed the data. TLS wrote the manuscript, with edits from all authors. All authors approved the manuscript.

## Acknowledgements

We would like to thank all Medical Entomology staff who were involved in mosquito surveillance and identification. We thank Nigel Beebe and Queensland Health for sample collection, and Kelly Richardson for assistance with library construction. Funding was provided by Australian Department of Health, Queensland Health, and the National Health and Medical Research Council (grant numbers 1132412, 1118640 and 1131932 (The HOT NORTH initiative)).

## Data Availability

Our data (mosquito .bam files and geographical locations) will be archived at the NCBI Sequence Read Archive by time of publication.

