## Supplemental Figures S1-S7 for "Genomic databanks and targeted assays help characterise domestic mosquito incursions"

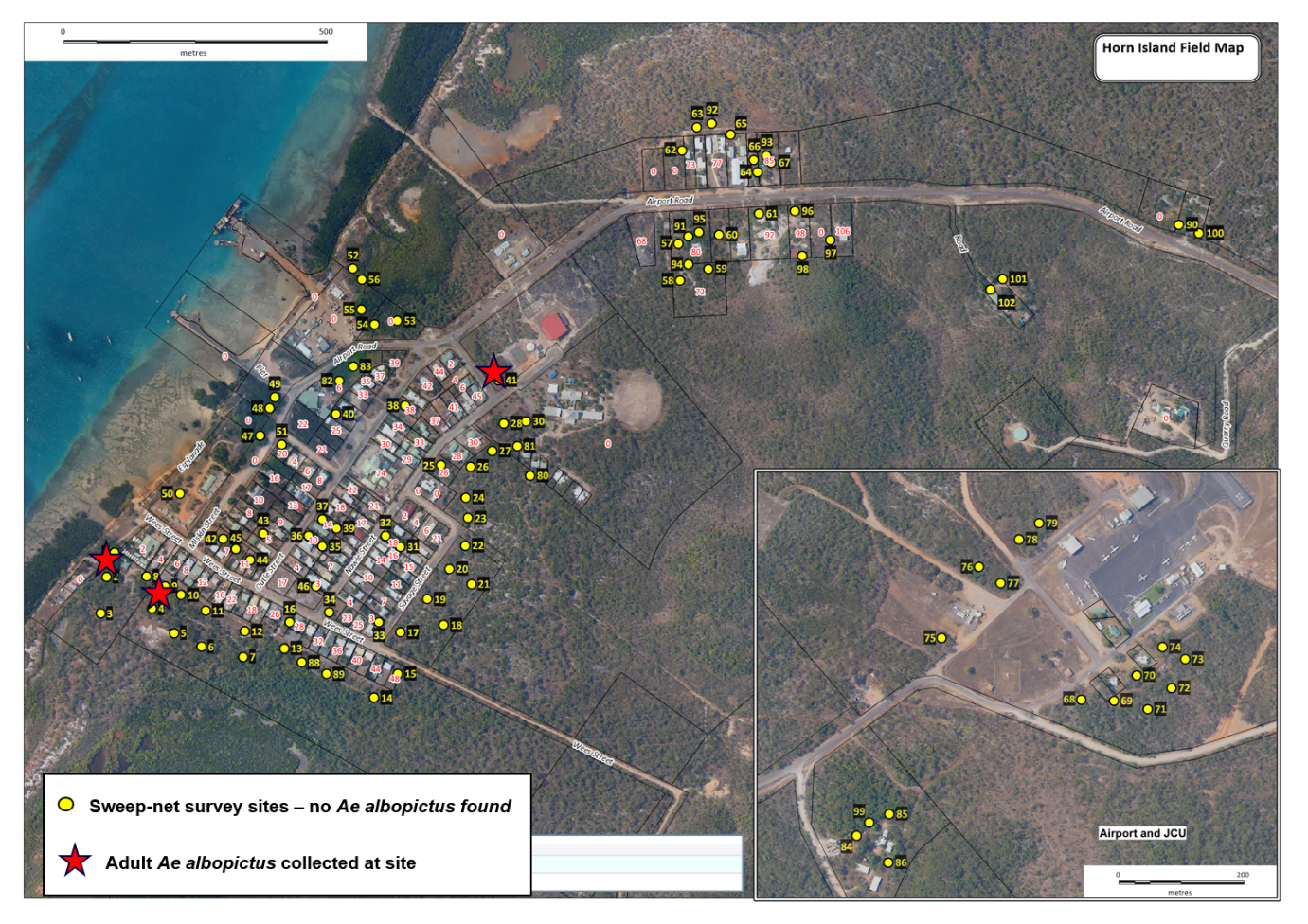

**Figure S1: Locations of the three *Aedes albopictus* collected on Horn Island in 2021 (red stars).** Yellow circles indicate sweep-net survey sites.

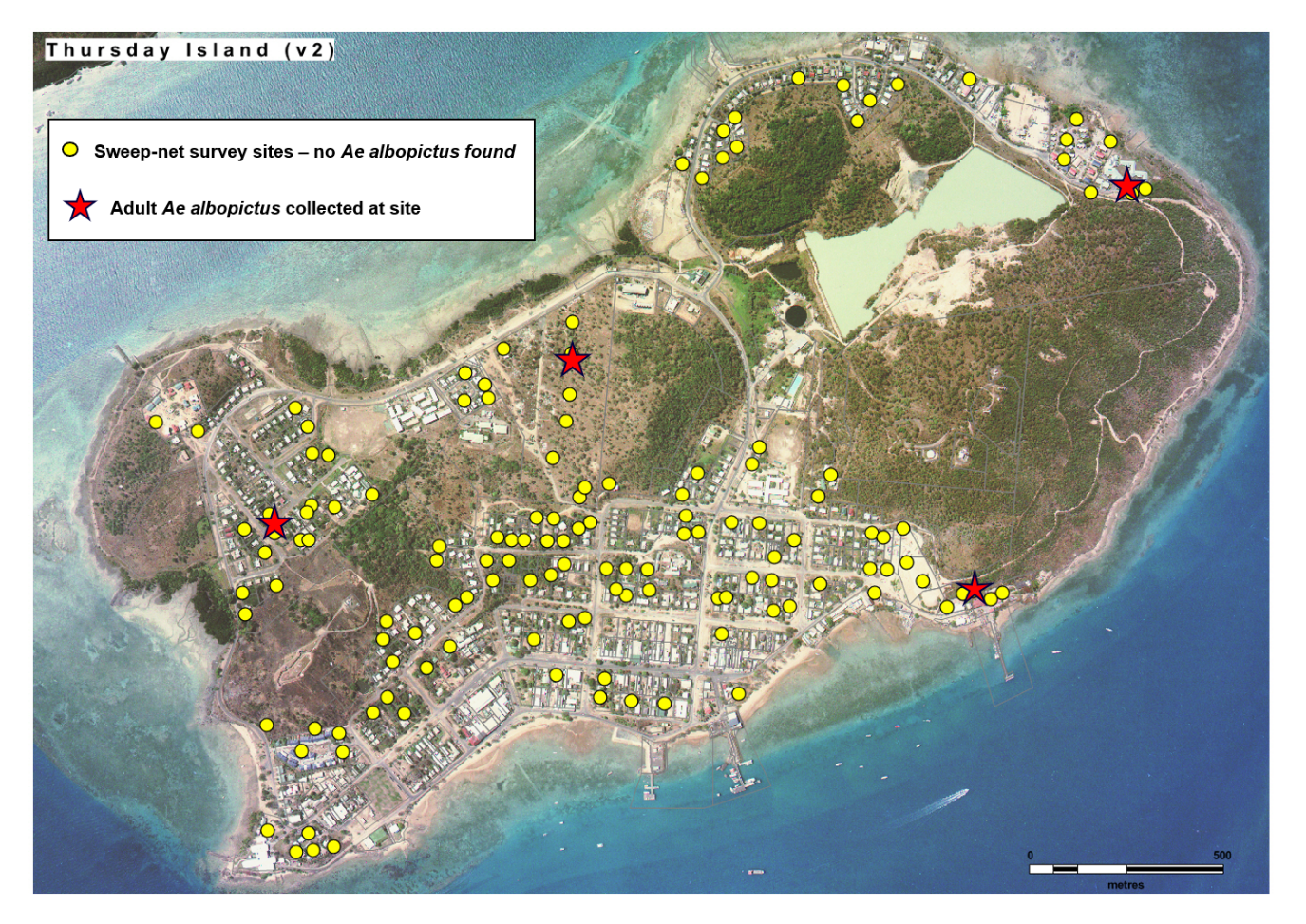

**Figure S2: Locations of the four *Aedes albopictus* collected on Thursday Island in 2021 (red stars).** Yellow circles indicate sweep-net survey sites.

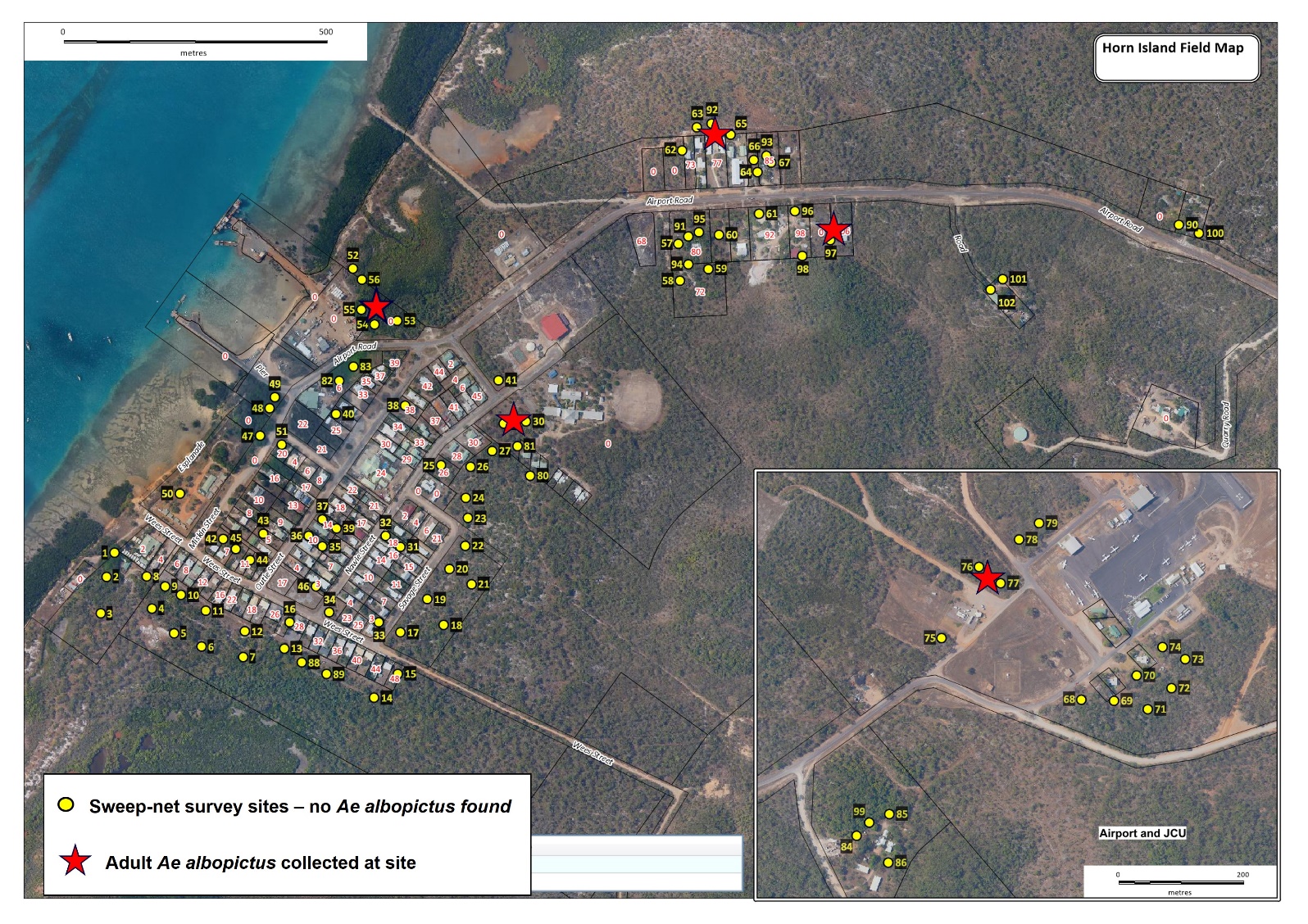
 **Figure S3: Locations of the five *Aedes albopictus* collected on Horn Island in 2022 (red stars).** Yellow circles indicate sweep-net survey sites.

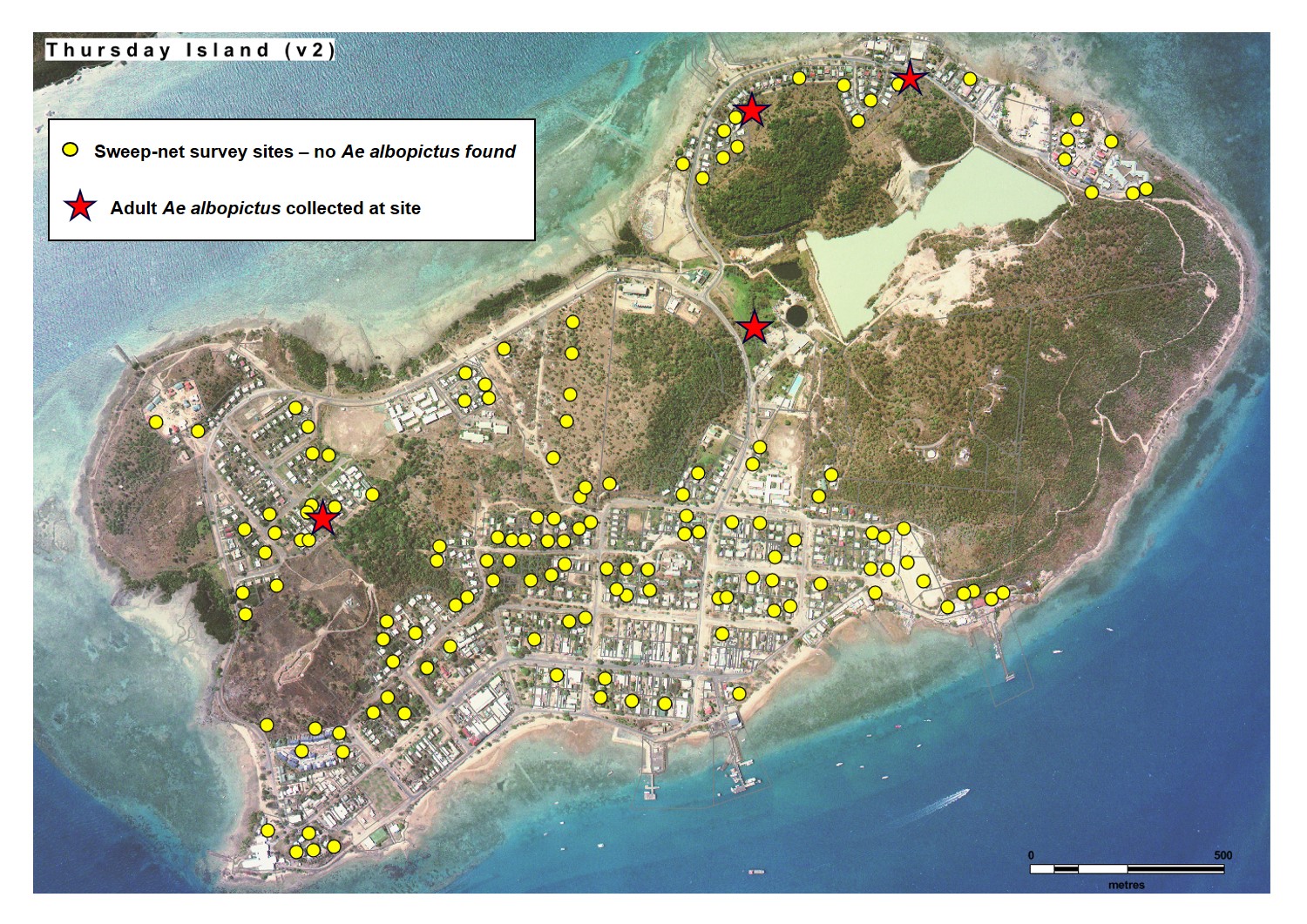
 **Figure S4: Locations of the four *Aedes albopictus* collected on Thursday Island in 2022 (red stars).** Yellow circles indicate sweep-net survey sites.

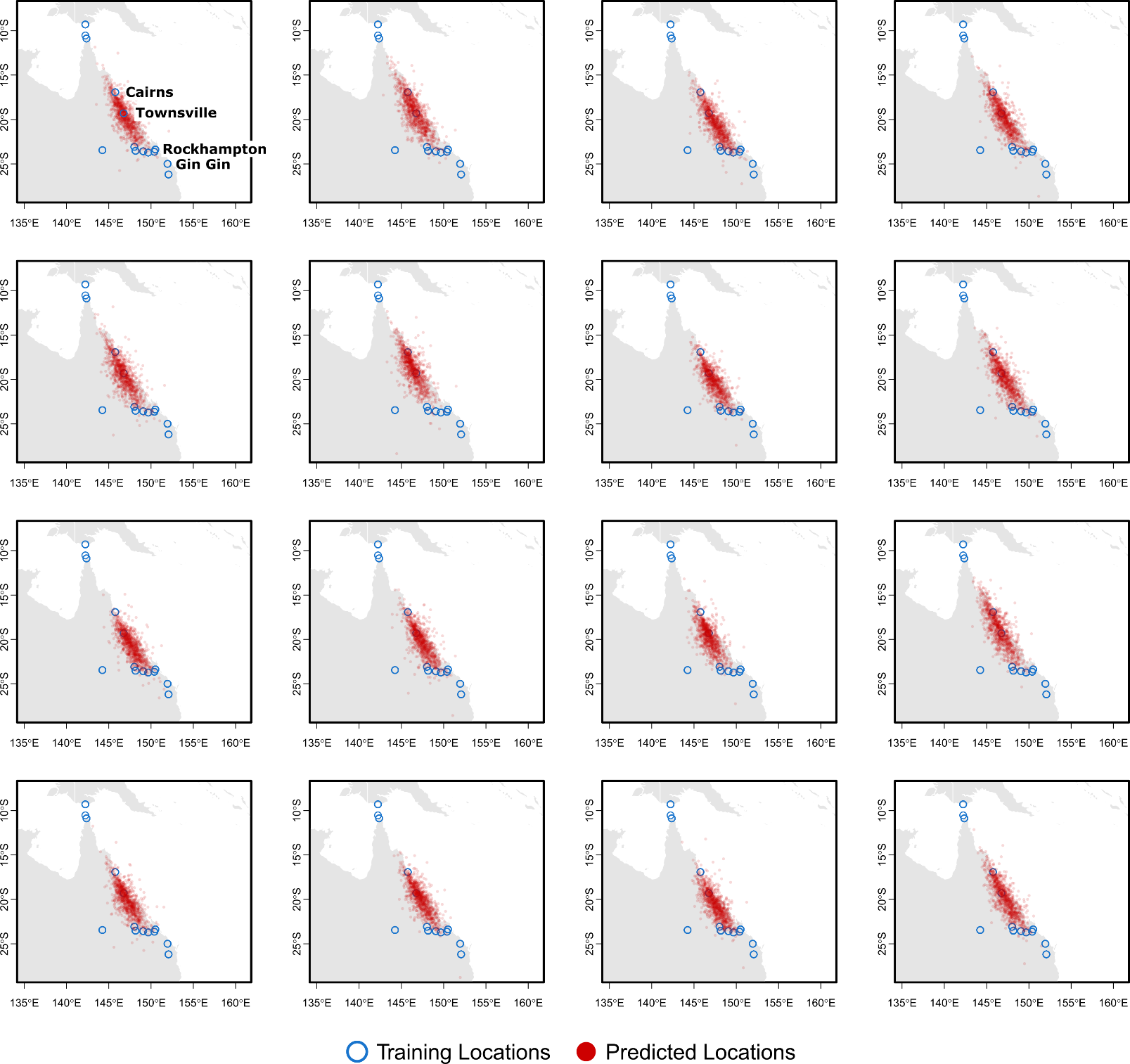

**Figure S5.** **Tracing of the 16 Tennant Creek individuals using the entire genomic databank.** Blue circles indicate reference sample locations, red circles show inferred origins from 1000 bootstrap replicate runs of Locator.

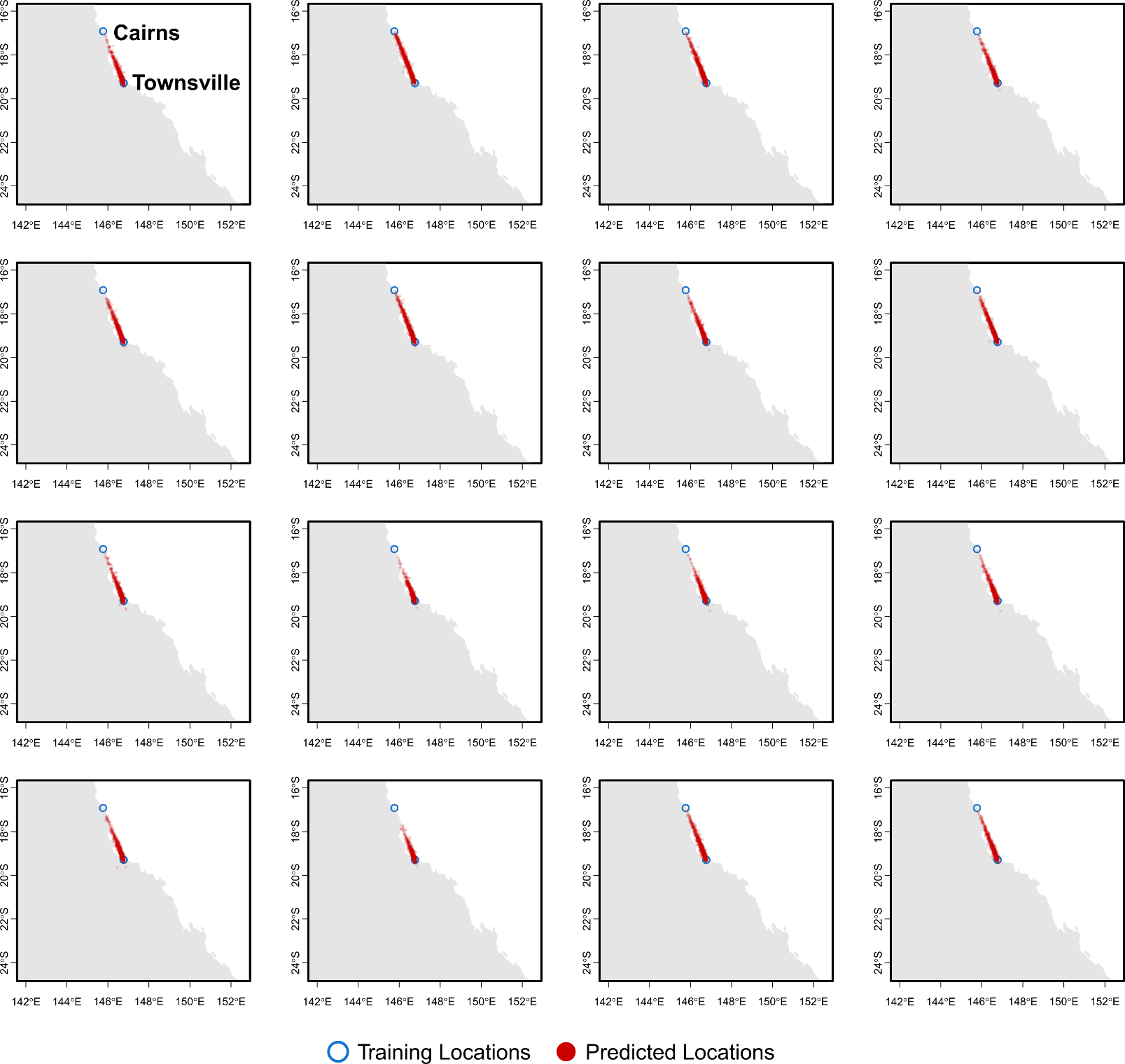

**Figure S6.** **Tracing of the 16 Tennant Creek individuals using only Cairns and Townsville.** Blue circles indicate reference sample locations, red circles show inferred origins from 1000 bootstrap replicate runs of Locator.

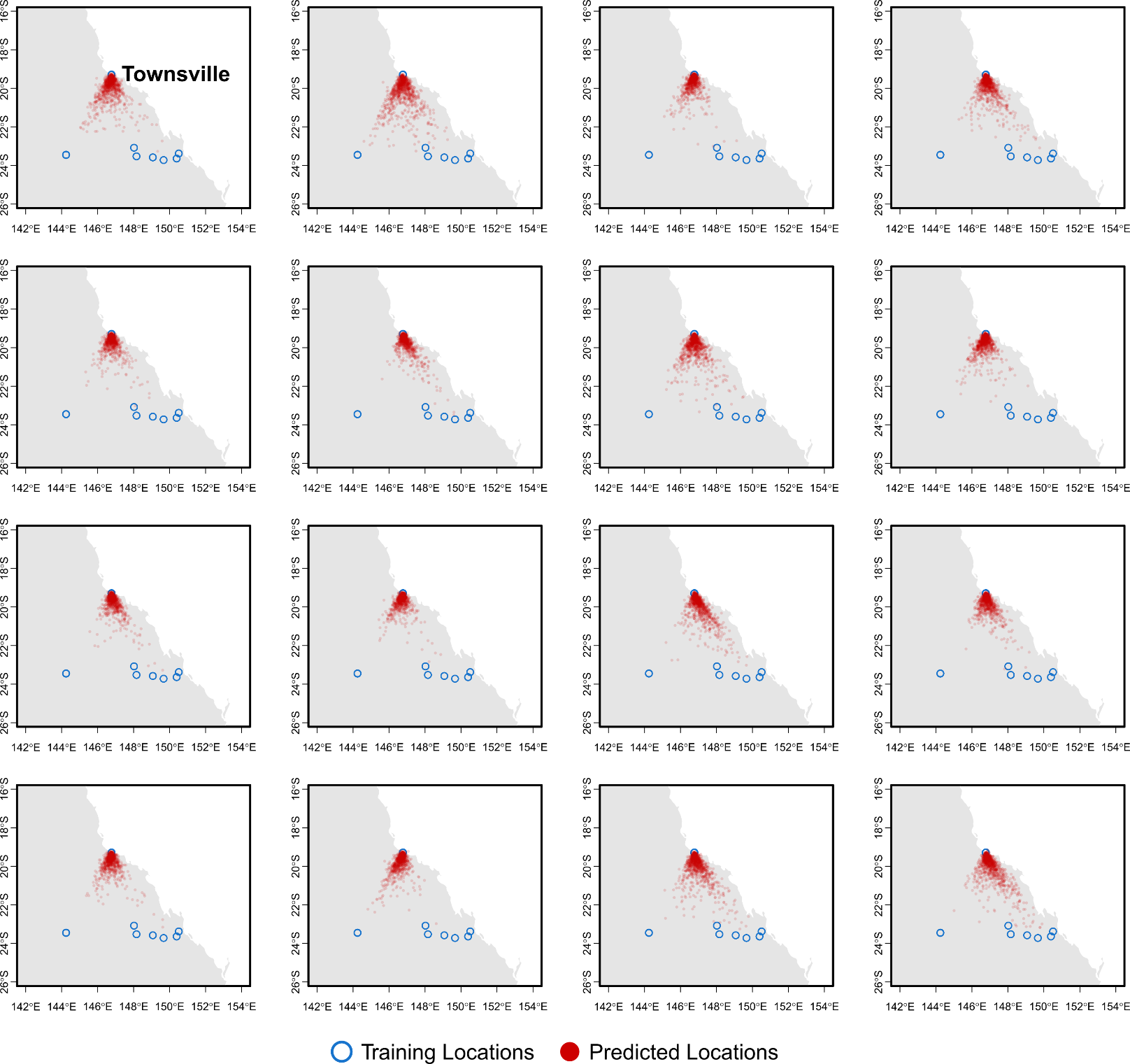

**Figure S7.** **Tracing of the 16 Tennant Creek individuals using only Townsville and nearby populations in southern Queensland.** Blue circles indicate reference sample locations, red circles show inferred origins from 1000 bootstrap replicate runs of Locator.
